# A method to assess the mitochondrial respiratory capacity of complexes I and II from frozen tissue using the Oroboros O2k-FluoRespirometer

**DOI:** 10.1101/2022.10.02.510534

**Authors:** Brad Ebanks, Nicoleta Moisoi, Lisa Chakrabarti

## Abstract

High-resolution respirometry methods allow for the assessment of oxygen consumption by the electron transfer systems within cells, tissue samples, and isolated mitochondrial preparations. As mitochondrial integrity is compromised by the process of cryopreservation, these methods have been limited to fresh samples. Here we present a simple method to assess the activity of mitochondria respiratory complexes I and II in previously cryopreserved murine skeletal muscle tissue homogenates, as well as previously frozen *D. melanogaster*, as a function of oxygen consumption.

## Introduction

As the field of biomedical research continues to expand throughout the 21^st^ century, the use of frozen tissue in research laboratories will continue to expand. This increases the accessibility of clinically relevant samples to researchers and institutions where fresh samples might not be available. Sample freezing has consistently proven to be effective when making assessments of genomes, transcriptomes and proteomes of a given sample^1^.

However, sample freezing is not always compatible with certain research interests, including mitochondrial respiration. Mitochondria are a fundamental organelle in eukaryotic organisms, where they are responsible for a wide range of biosynthetic and metabolic processes. This includes haem biosynthesis^2^, calcium signalling^3^, and apoptosis regulation^4^. However, they are most widely studied for the capacity to produce the biological energy currency of ATP, through the coupling of oxidative phosphorylation with the electron transfer system.

The electron transfer system is composed of a series of enzyme complexes in the inner mitochondrial membrane that oxidise a range of metabolites, including NADH derived from the TCA cycle^5^, fatty acids via beta-oxidation^6^, succinate^7^, proline^8^, and glycerol-3-phosphate in mammals^9^. Electrons derived from these species are funnelled into the Q-junction (co-enzyme Q), before the sequential movement through complex III, cytochrome c and finally respiratory complex IV, where they reduce molecular oxygen in a system that has been well-reviewed^10^. As a part of this process, complexes I, III, and IV pump protons across the inner mitochondrial membrane to generate the proton motive force that drives ATP synthesis via ATP synthase^11^.

Classical studies by Chance and Williams in the 1950s used platinum microelectrode chemistry to assess oxygen consumption, with simultaneous spectrophotometric assays to assays ATP production via DNPH oxidation^12^. The current leading instruments for assessing mitochondrial oxygen consumption are based on either fluorometric measurement of oxygen (SeaHorse)^13,14^, or Clark electrodes within polarographic oxygen sensors (Oroboros Oxygraph-O2k)^15^. In particular the Oroboros Oxygraph-O2k allows for dynamic high-resolution respirometry assessments across tissues, cells, and mitochondrial isolates in response to titrations of substrates, uncouplers, and inhibitors of mitochondrial respiratory function (SUIT protocols)^16^.

One limiting factor of existing SUIT protocols for the Oroboros Oxygraph-O2k, and respirometry assessments made with other instruments, is the inability to assess the respiratory profile or frozen tissue samples. This is due to the inactivation of the TCA cycle and the rupturing of the mitochondrial outer membrane during the process of sample freezing^17–19^. However, there are established enzyme assays that can measure the activity of mitochondrial enzyme complexes^20^.

Recent studies have described methods to assess mitochondrial oxygen consumption in cryopreserved samples^21^, alongside new respirometry protocols in different instruments^22^. Therefore we sought to establish a method to measure the activity of mitochondrial respiratory complexes I and II as a function of oxygen consumption, in the Oroboros Oxygraph-O2k. NADH is directly oxidised by complex I, and succinate is commonly used in respirometry to assess complex II activity, so we tested these substrates in homogenates of previously frozen mouse tissue and *Drosophila*.

## Methods

### Mouse husbandry

Skeletal muscle sourced from 71-week-old female C57BL/6J mice was sourced from Charles River. Samples were snap frozen and stored at −80°C.

Animals were bred and housed in accordance with strict Home Office stipulated conditions. The overall programme of work (in respect to the original UK Home Office Project Licence application) is reviewed by the Animal Welfare and Ethical Review Body at the University of Nottingham and then scrutinised by the UK Home Office Inspectorate before approval by the Secretary of State. Individual study protocols link to the overarching Home Office Project Licence and are made available to the Named Animal Care and Welfare Officer, the Named Veterinary Surgeon (both are members of the AWERB), the animal care staff and the research group. The Project Licence Number for the breeding and maintenance of this genetically altered line of mice is PPL 40/3576. The mice are typically group housed and maintained within solid floor cages containing bedding and nesting material with additional environmental enrichment including chew blocks and hiding tubes. Cages are Individually Ventilated Cage Units within a barrier SPF unit to maintain biosecurity. Animals are checked daily by a competent and trained animal technician. Any animal giving cause for concern such as subdued behaviour, staring coat, loss of weight or loss of condition will be humanely killed using a Home Office approved Schedule 1 method of killing.

### Drosophila husbandry

*Drosophila melanogaster* strain w1118 (males) were used in this study, housed in glass vials Fly food (Quick Mix Medium, Blades Biological) was added to the vial, to a depth of 1 cm, and 3 mL of distilled water was added; it was left for one minute, and then a small sprinkle of yeast was added. The *Drosophila* were maintained at 25°C, and the food was rehydrated with 150 uL of H2O every 24 hours. Flies were either frozen at −20 or −80. The study was reviewed and approved by the University of Nottingham SVMS local area ethics committee (#3091 200203 10 February 2020).

### Skeletal muscle sample preparation

5, 10, and 25 mg of skeletal muscle tissue from 71-week-old female mice was mechanically homogenised in 300 μL of MiRO5 buffer (Oroboros Instruments; 0.5 mM EGTA, 3 mM MgCl2, 60 mM lactobionic acid, 20 mM taurine, 10 mMKH2PO4, 20 mM HEPES, 110 mM D-sucrose, 1 g/L BSA, pH 7.1). The homogenate was spun down at 850 g (10 mins, 4°C) to remove the insoluble fraction. The subsequent lysate was added to 2ml volume chambers in the Oroboros O2k-FluoRespirometer. Technical replicates were used for this study (N=5).

### Drosophila sample preparation

Five flies were mechanically homogenised in 500 μL of MiR05 buffer. The homogenate was spun at 850 g (10 mins, 4°C) to clear insoluble material. The supernatant was added to the O2k-FluoRespirometer. Biological replicates were used for this study, with samples frozen at −20 (n=3) or −80 (n=6) used in this study.

### High-resolution respirometry

A substrate, uncoupler, inhibitor, titration (SUIT) protocol was used to assess the uncoupled mitochondrial oxygen consumption capacity^16^. Briefly, 5 μL of cytochrome c was titrated into each chamber and a baseline value was reached, this was followed by a titration of 10 μL of NADH (10 mM) and the peak specific flux value was marked, then 20 μL of succinate (1 M), 1 μL of rotenone (1 mM), and 1 μL of antimycin A (5 mM) which is used for background correction.

Data analyses were performed in GraphPad Prism version 9.3.1.

## Results

### The mitochondrial respiratory capacity of frozen mouse skeletal muscle homogenate

Different masses of skeletal muscle tissue were homogenised before being assessed for their mitochondrial respiratory capacity in response to NADH and succinate titrations (**Figure 1**). All tissues masses that we assessed (5 mg, 10 mg, 25 mg) showed oxygen consumption in response to substrate titration, and the 5 mg tissue sample lysate produced the strongest oxygen consumption signal (**Table 1**).

**Figure 1.**
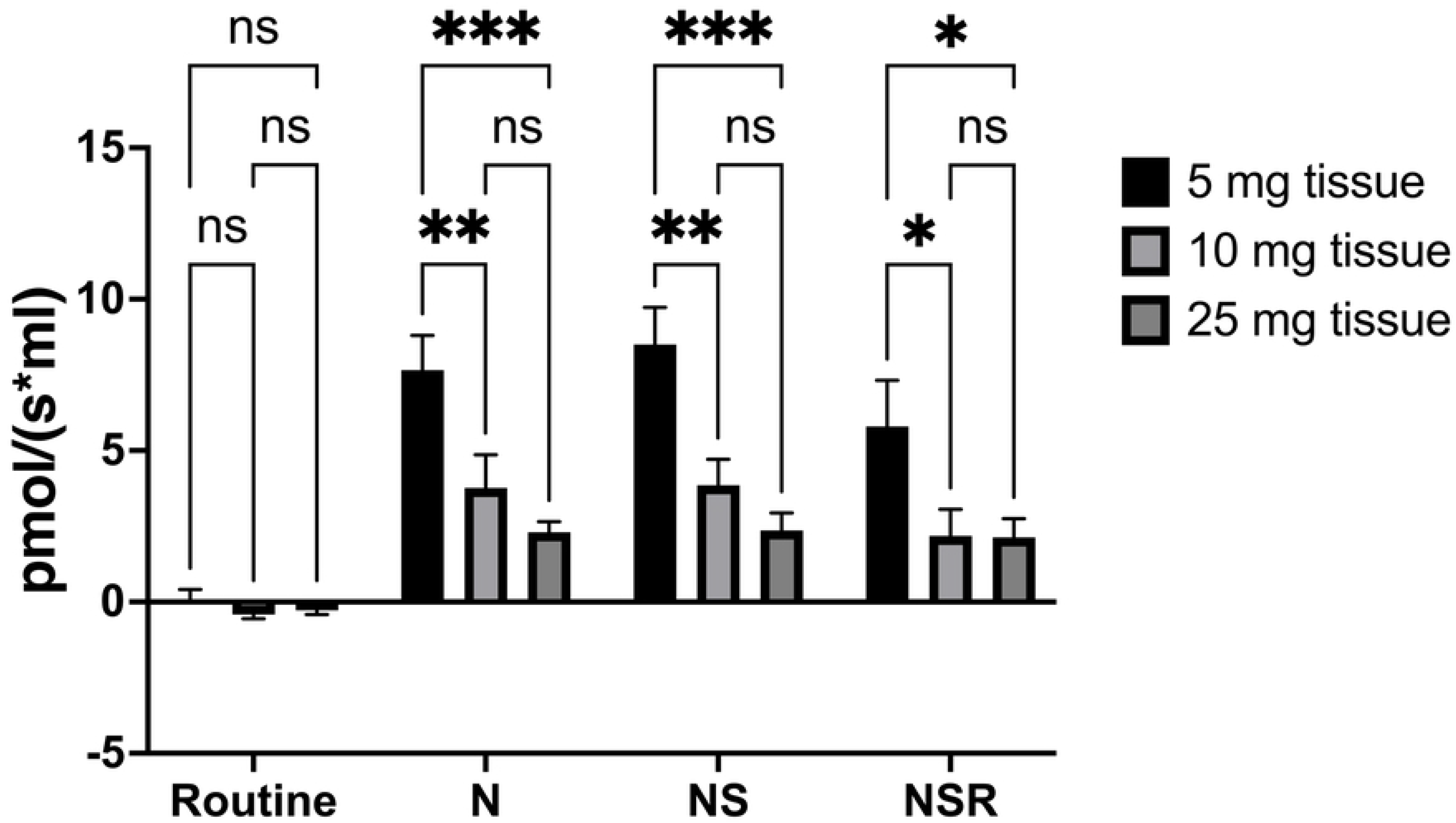
Complex I and complex II-linked mitochondrial oxygen consumption of frozen mouse skeletal muscle. Mitochondrial respiration was stimulated in either 5,10, or 25 mg mouse skeletal muscle tissue homogenates with titrations of NADH (N) and succinate (S), before being inhibited by rotenone (R), and antimycin A for background correction. Technical replicates, N = 5, 2-way ANOVA, Turkey’s multiple comparison test, * P = 0.033, ** P = 0.002, *** P < 0.001.

**Table 1.** Specific flux values of mouse skeletal muscle oxygen consumption. Mean specific flux values (pmol/(s*mL)) in 5 mg, 10 mg, and 25 mg lysates of skeletal mouse muscle tissue in response to sequential titrations of substrates (NADH, succinate) and inhibitors (rotenone). N=5, (SEM).

| Specific flux of substrate and inhibitor titrations (pmol/(s*mL)) | Mouse skeletal muscle |  |  |
| --- | --- | --- | --- |
|  | 5 mg | 10 mg | 25 mg |
| **Routine** | 0.016 (0.396) | -0.415 (0.136) | -0.281 (0.135) |
| **NADH** | 7.659 (1.139) | 3.769 (1.097) | 2.304 (0.345) |
| **NADH, succinate** | 8.501 (1.228) | 3.852 (0.863) | 2.358 (0.579) |
| **NADH, succinate, rotenone** | 5.795 (1.527) | 2.181 (0.876) | 2.13 (0.618) |

### The mitochondrial respiratory capacity of frozen Drosophila (flies) after homogenisation

Using the same protocol, we assessed the activity of complex I and complex II as a function of oxygen consumption in *D. melanogaster* that had previously been frozen at either −20°C or −80°C (**Figure 2**). We report that oxygen consumption in *D. melanogaster* frozen at −20°C is limited for titrations of both NADH and succinate (**Table 2**), but a strong signal is detected in *D. melanogaster* that had previously been frozen at −80°C.

**Figure 2.**
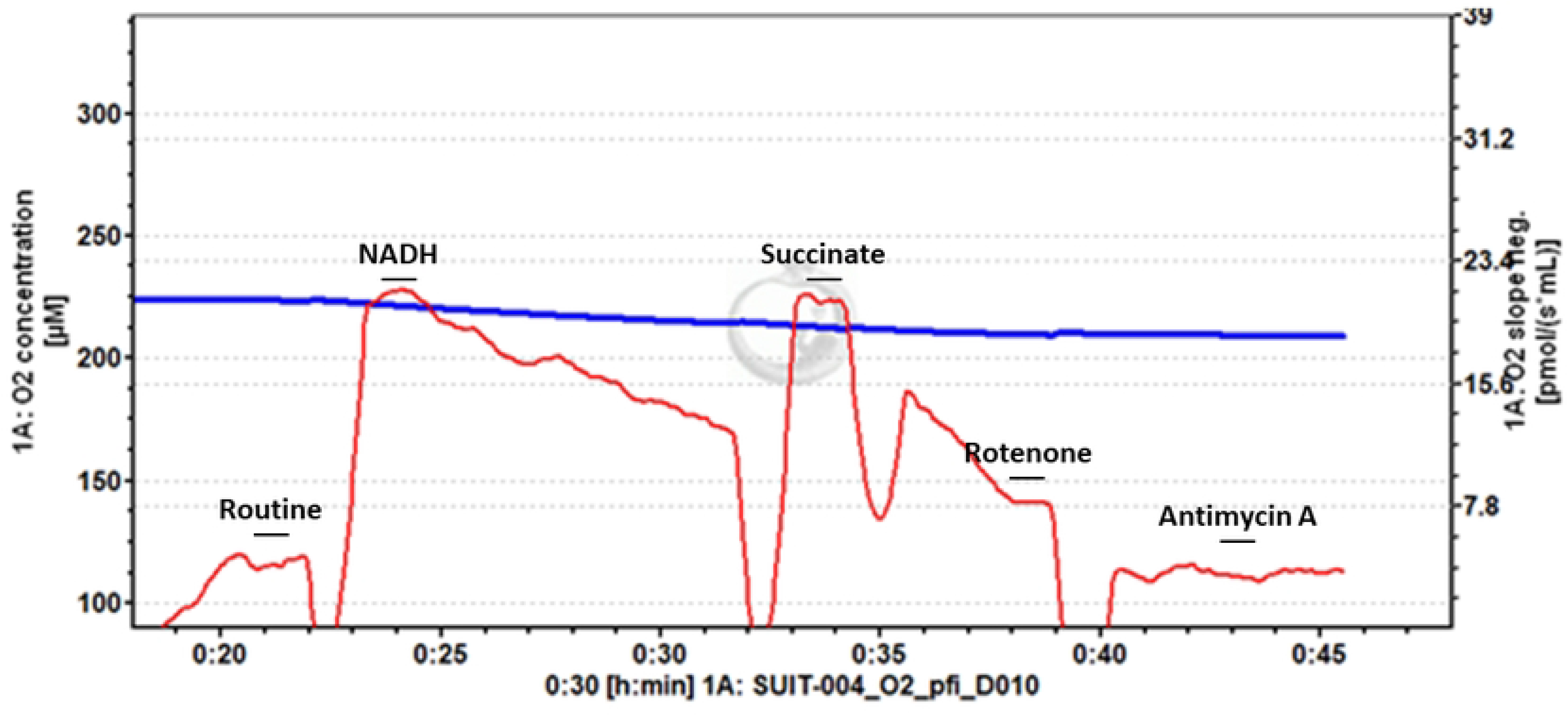
Example trace of frozen mouse skeletal muscle. Stepwise titrations of NADH and succinate stimulate oxygen flux, which is inhibited by rotenone and antimycin A titrations. Titration of NADH leads to a peak of oxygen flux, before progressive decrease.

**Figure 3.**
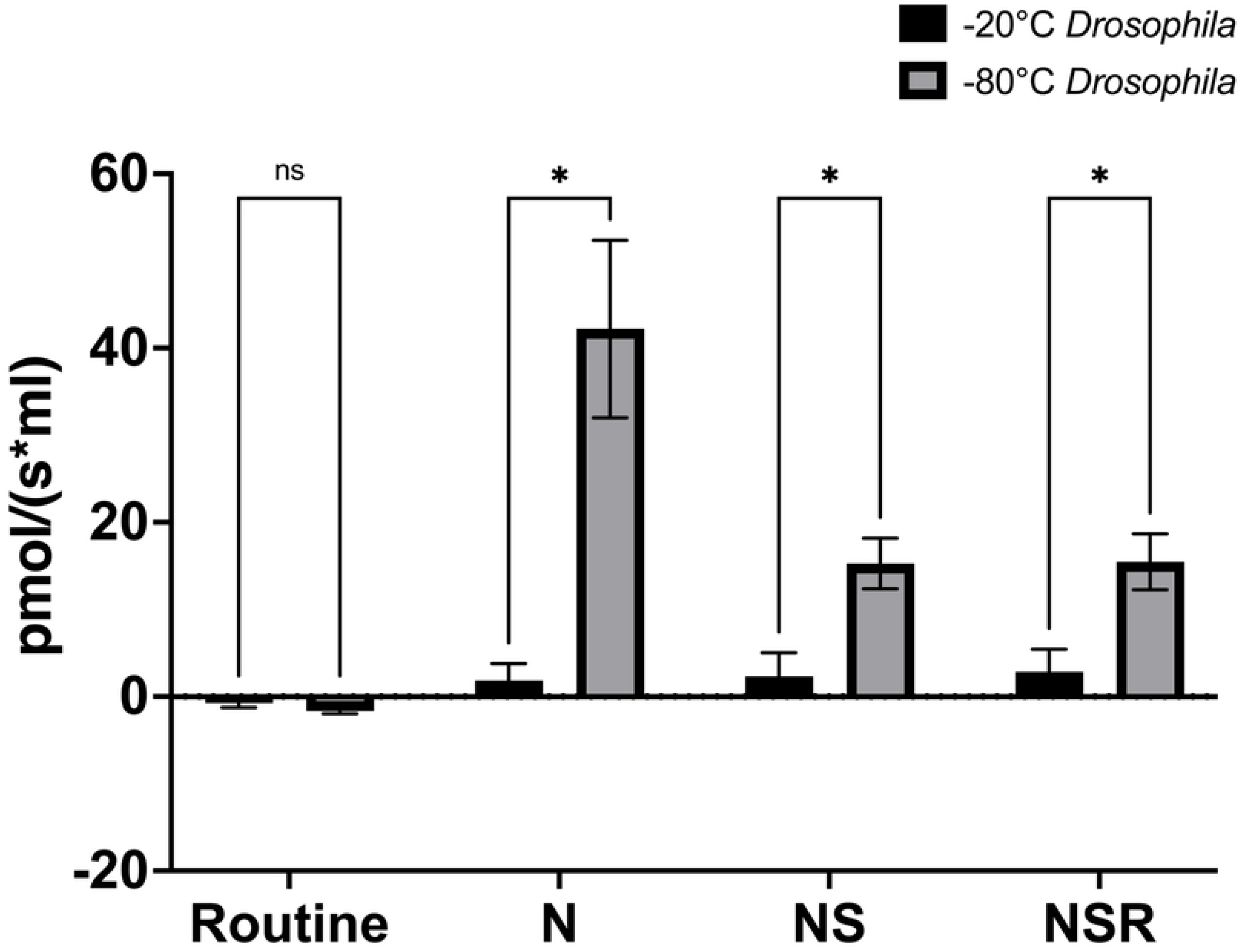
Complex I and complex II-linked mitochondrial oxygen consumption of D. melanogaster frozen at −20 °C and −80 °C. The specific oxygen flux (pmol/(s*mL)) of D. melanogaster was assessed in samples that had been frozen at either −20 °C (black) or −80 °C (grey) in response to stepwise titrations of NADH (N), succinate (S), and rotenone (R), followed by antimycin A for background correction. D melanogaster −20 °C (N=3), D. melanogaster −80 °C (N=6). Error bars = SEM, * p < 0.05.

**Figure 4.**
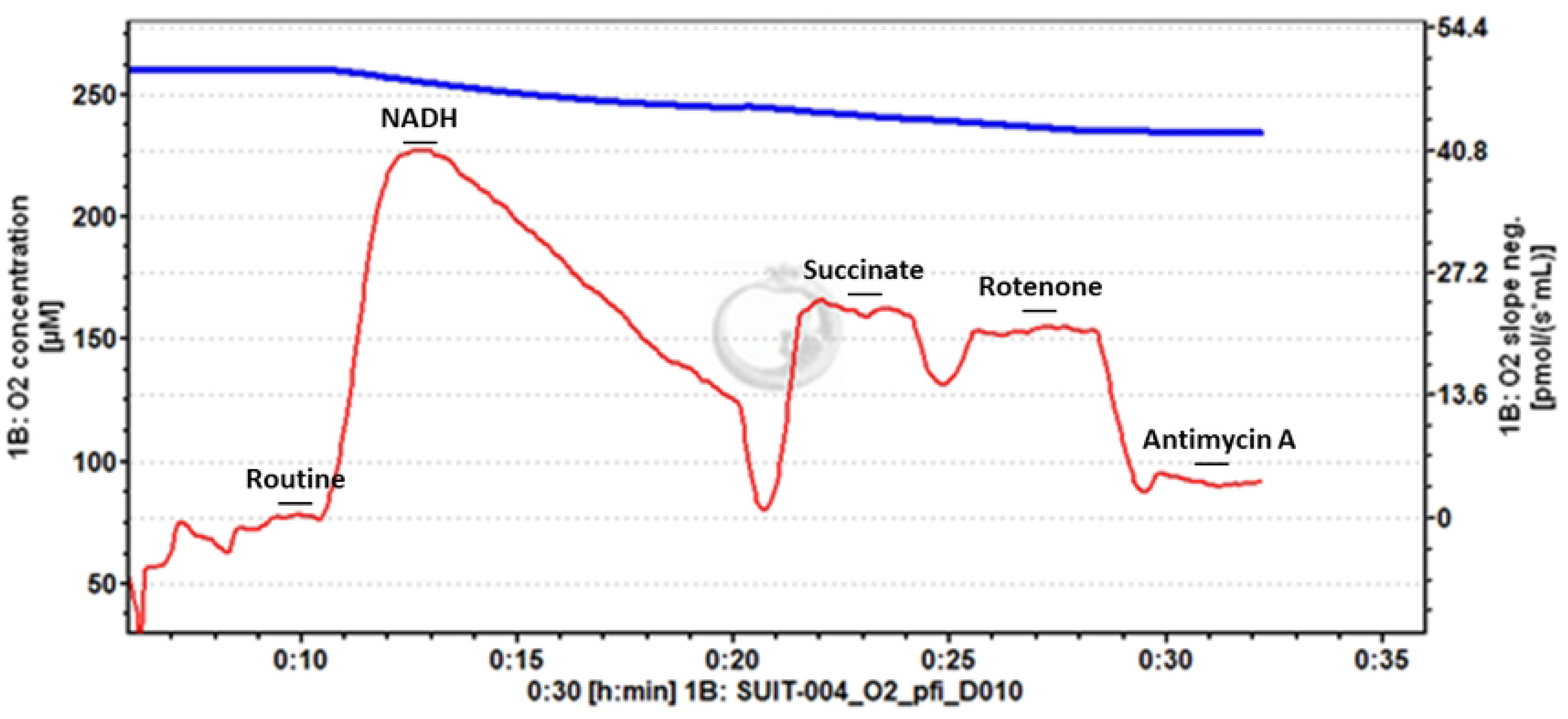
Example trace of −80°C frozen Drosophila. Stepwise titration of NADH and succinate stimulate oxygen flux, that is then inhibited by titrations of rotenone and antimycin A. Titration of NADH leads to a peak of oxygen flux, before progressive decrease.

**Table 2.** Specific flux values of oxygen consumption from previously frozen D. melanogaster. Mean specific flux values (pmol/(s*mL)) in homogenates of D. melanogaster previously frozen at −20°C and −80°C, in response to sequential titrations of substrates (NADH, succinate) and inhibitors (rotenone). D melanogaster −20°C (N=3), D. melanogaster −80°C (N=6), (SEM).

| **Specific flux of substrate and inhibitor titrations (pmol/(s\*mL))** | ***D. melanogaster*** |  |
| --- | --- | --- |
|  | **-20** | **-80** |
| **Routine** | -0.745<br>(0.499) | -1.658<br>(0.310) |
| **NADH** | 1.847 (1.928) | 42.19<br>(10.176) |
| **NADH, succinate** | 2.334 (2.679) | 15.26 (2.915) |
| **NADH, succinate, rotenone** | 2.825 (2.608) | 15.48 (3.205) |

## Discussion

We have demonstrated a simple method to assess the activity of mitochondrial electron transport complexes I and II as a function of mitochondrial oxygen consumption using the Oroboros Oxygraph-O2k. We demonstrated the feasibility of this protocol using lysates of different masses from mouse skeletal muscle, as well as comparing the values between *D. melanogaster* that had previously been frozen at either −20°C or −80°C.

We report that when the 2ml volume chambers are used the smallest mass tested of mouse skeletal muscle homogenate, 5 mg, gives the strongest signals in response to the substrate and inhibitor titrations (**Figure 1**). The observation of a stronger signal being observed for the smallest mass of tissue could be due to an excess of tissue obscuring the oxygen flux detection by the instrument, and warrants additional investigation. For the *D. melanogaster*, those that had been frozen at −80°C gave a significantly stronger signal than those frozen at −20°C (**Figure 2**), which is understandable due to the colder temperatures better maintaining the structural integrity of the relevant biomolecules.

While freezing of samples is known to compromise the integrity of the outer mitochondrial membrane^23,24^, thus preventing the study of coupled respiratory capacity and oxidative phosphorylation^25^, this study shows that the electron transfer system is maintained in such a way that the activity of constituent enzymes can be assessed as a function of oxygen consumption by the system. This is in contrast with static assays which measure the specific activity of the single enzyme in isolation^20,26^.

Previous studies, including in the Oroboros Oxygraph-O2k^27^, have presented methods to assess the oxygen consumption of the electron transfer system from cryopreserved samples^21,22^. Through consideration of these studies, we have developed the methods reported in this study.

We also believe that this protocol is highly useful for assessing the activities of respiratory complexes in samples that have previously been archived at −80°C, including valuable clinical samples. It may also be considered for use by research groups that are based in laboratories and institutions where access to high-quality fresh samples is a particular logistical challenge.

Further work using the principles outlined in this pilot study could be undertaken to explore the feasibility of assessing the activity of other electron transfer system enzymes, including proline dehydrogenase, the electron-transferring flavoprotein complex (cETF), and the mitochondrial glycerol-3- phosphate dehydrogenase in mammalian systems.

## Author Contributions

B.E. performed the experiments, completed the data analysis, and wrote the manuscript. N.M. bred and provided *D. melanogaster*. L.C. directed the research, supervised experiments, provided reagents, and prepared the manuscript.

